# Elucidating interplay of speed and accuracy in biological error correction

**DOI:** 10.1101/102608

**Authors:** Kinshuk Banerjee, Anatoly B. Kolomeisky, Oleg A. Igoshin

## Abstract

One of the most fascinating features of biological systems is the ability to sustain high accuracy of all major cellular processes despite the stochastic nature of underlying chemical processes. It is widely believed that such low errors are the result of the error correcting mechanism known as kinetic proofreading. However, it is usually argued that enhancing the accuracy should result in slowing down the process leading to so-called speed-accuracy trade-off. We developed a discrete-state stochastic framework that allowed us to investigate the mechanisms of the proofreading using the method of first-passage processes. With this framework, we simultaneously analyzed speed and accuracy of the two fundamental biological processes, DNA replication and tRNA selection during the translation. The results indicate that speed-accuracy trade-off is not always observed. However, when the trade-off is present, the biological systems tend to optimize the speed rather than the accuracy of the processes, as long as the error level is tolerable. Additional constraints due to the energetic cost of proofreading also play a role in the error correcting process. Our theoretical findings provide a new microscopic picture of how complex biological processes are able to function so fast with a high accuracy.

---

**B**iological systems exhibit remarkable accuracy in selecting the right substrate from the pool of chemically similar molecules. This is common to all fundamental biological processes such as DNA replication, RNA transcription and protein translation [1]. The level of fidelity in various stages of genetic information flow depends on their relative importance in sustaining system stability. DNA replication is thought to be the most accurate process with an error rate, *η* ~ 10^−8^ −10^−10^ [2, 3] *i.e.*, only 1 out of 10^8^ − 10^10^ incorporated nucleotides is mismatched. RNA transcription (*η* ~ 10^−4^−10^−5^) and protein translation (*η* ~ 10^−3^ −10^−4^) processes are also quite accurate but to a somewhat lower degree [4, 5]. Failure to maintain such accuracy adversely affects cell viability and survival. For example, mutations affecting the fidelity in translation increase the amount of unfolded proteins leading to apoptosis [6] and to erroneous replication of genetic material [7].

Initially it was unclear how the small differences in equilibrium binding stability of structurally similar substrates can allow such a high degree of discrimination [8]. Then, an explanation was provided, independently by Hopfield and Ninio [9, 10], who proposed an error-correction mechanism called kinetic proofreading (KPR). KPR allows enzymes to utilize the free energy difference between right and wrong substrates multiple times using additional steps [9]. This amplifies the small energetic discrimination and results in a lower error compared to that in chemical equilibrium. However, such processes require significant energy consumption [9]. To this end, enzymes employ some energy-rich molecules, like ATP, to provide for the necessary driving [11, 12]. The mechanism was experimentally verified later in different biological systems [13–17]. Several recent studies generalized it to more complex networks and found analogies between proofreading and other phenomena such as microtubule growth [18] or bacterial chemotaxis [19]. These results broaden the concept of KPR and show that such chemically-driven regulatory mechanisms are widely present.

Cells must process genetic information not just accurately but also sufficiently rapidly. Proofreading enhances the accuracy by resetting the system to its initial configuration without progressing to product state [9]. The completion time of the reaction is, thus, expected to increase. Hence, there could be a compromise, or trade-off, between accuracy and speed of the process [20]. The understanding on this trade-off is mainly based on the Michaelis-Menten (MM) description of specificity [21, 22]. These studies indicate that, the minimum-possible error is achieved at vanishingly-low catalytic rate, i.e. when the process is the slowest [9, 21]. In contrast, biological polymerization reactions must occur reasonably fast [15, 23]. A recent study demonstrated a new speed-accuracy regime in the KPR model by modifying the catalytic rate [18]. In this regime, a large gain in speed comes with a relatively small loss in accuracy. The authors suggested that biological systems may employ this regime [18]. For example, in tRNA selection process, fast GTP hydrolysis step speeds up protein synthesis but prevents maximal possible selectivity of the initial tRNA-ribosome binding step [21, 24].

### Significance Statement

Biological processes are unique by showing a remarkable level of accuracy in discriminating between similar molecules. This is attributed to an error-correcting mechanism known as kinetic proofreading. It is widely believed that the enhancement of the accuracy in biological processes always slows down them. Our theoretical study reveals that such trade-offs might not always happen. By analyzing the fundamental processes of DNA replication and protein translation, we established that these systems maximize speed rather than accuracy with additional energetic constraints. Our findings provide a microscopic picture of how complex biological processes can be accomplished so quickly with minimal errors.

Despite the number of studies, a clear quantitative picture of how the balance between speed and accuracy is tuned is lacking. Several current models of proofreading, still mainly focus on the initial stages of substrate selection [22, 25, 26] or assume disparity of rate constants of only a few type of steps [18, 19]. In contrast, experimental data show that biological systems have different rates for the right and wrong substrates for each step of the network [4, 14, 15]. Therefore, by distributing the discrimination of the reaction rates over the whole network, biological systems might be able to achieve better balance between speed and accuracy. As a consequence, regimes with no trade-off may also arise. Moreover, proofreading steps come with an extra energy cost to gain higher accuracy [11] but the role of this cost in the trade-off is not apparent. Therefore, to understand the fundamental mechanisms of proofreading in real biological systems, one needs to answer the following questions: (i) How does the system set its priorities when choosing between accuracy and speed, two seemingly opposite objectives? (ii) Can speed and accuracy change in the same direction or, in other words, is there always a trade-off? (iii) How does the extra energy expenditure due to KPR affect the speed-accuracy optimization?

Here we focus on the role of reaction kinetics in governing the speed-accuracy trade-off. To this end, we develop a generalized framework to study one-loop KPR networks, assuming distinct rate constants for every step of the right and wrong (W) pathways. Based on this approach, we model the overall selection of correct substrate over the incorrect one as a first-passage problem to obtain a full dynamic description of the process [27, 28]. This general framework is applied to two important examples, namely, DNA replication by T7 DNA polymerase (DNAP) [3, 14] and protein synthesis by *E.coli* ribosome [22, 29] (Fig. 1(A,B)). Starting from the experimentally measured rate constants for each system we vary their values to analyze the resulting changes in speed and accuracy and to assess the trade-off. The role played by the extra energy consumption or cost of proofreading [11] is also investigated. By comparing the behavior of the two systems, we search for general properties of biological error correction.

**Fig. 1.**
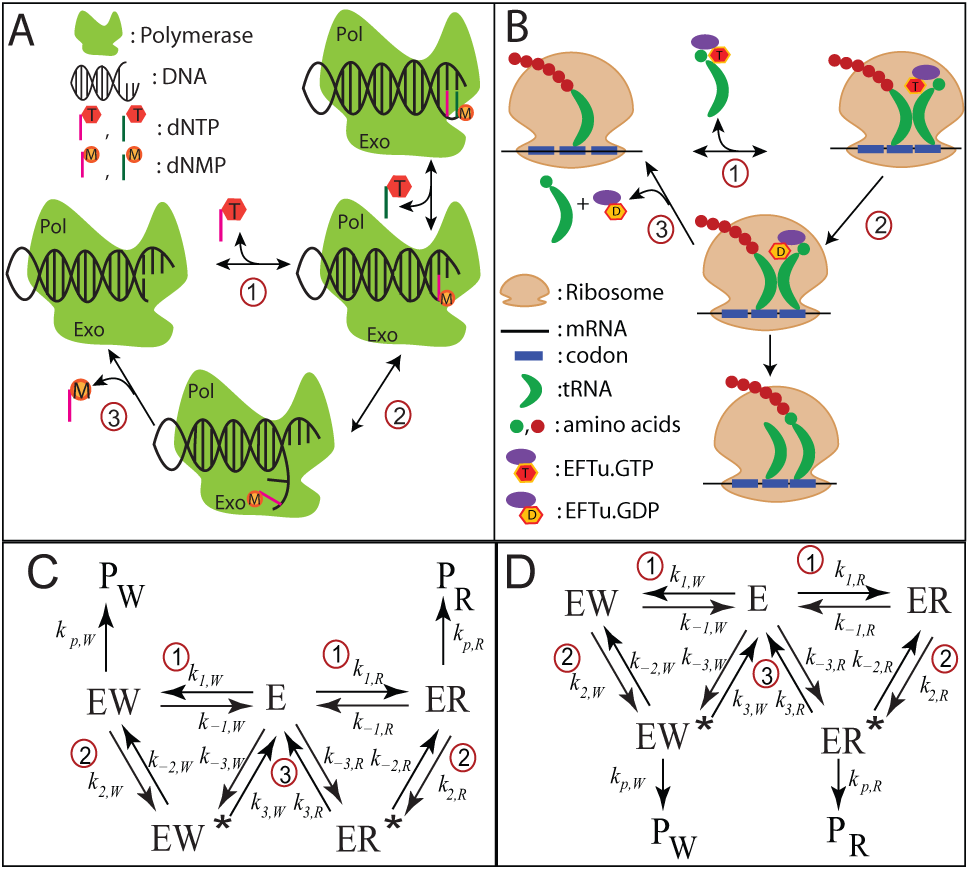
Schematic representation of proofreading networks for (A) DNA replication byT7 DNA polymerase (DNAP) enzyme and (B) aminoacyl(aa)-tRNA selection by *E.coli* ribosome during translation. Corresponding chemical networks are shown for replication and (D) translation. Reaction steps comprising the cycles are labeled 1-3. Rate constants of each step are denoted by *k*_±*i,R/W*_, *i* = 1, 2, 3; subscript R or W indicates right (R) or wrong (W) pathways. The rate constants of the steps leading to product (end) states are labeled as *k*_p,R*/*W_. The translation network in is related to the replication network in (C) by the following transformation of rate constant indices: 
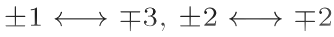
. The steps involved in each case are, of course, different. For details, see text.

## Methods

### Proofreading networks of replication and translation

DNA replication as well as protein synthesis employ nucleotide complementarity to select the cognate substrate over others. During replication, dNTP molecules complementary to the DNA template are chosen. Similarly, during protein synthesis, aminoacyl(aa)-tRNAs are picked by ribosome based on the complementarity of their anti-codon to the mRNA codon. Wrong substrates that bind initially can be removed by error-correction proofreading mechanisms. Kinetic experiments coupled with modeling revealed a lot of mechanistic details about both the processes [3, 14, 15, 24]. The schemes depicted in Fig. 1 represent the key steps to understand the KPR in these networks.

The schemes in Fig. 1(A,C) are for DNA replication [3, 14]. E denotes the T7 DNAP enzyme in complex with a DNA primer-template. The right and wrong substrates are correct and incorrect base-paired dNTP molecules, respectively. Step generates enzyme-DNA complexes ER(or EW) with the primer elongated by one nucleotide. Addition of another correct nucleotide to ER (EW) gives rise to P_R_ (P_W_). ER^*^ and EW^*^ complexes denote the primer shifted to the exonuclease site (Exo) from the polymerase site (Pol) of DNAP. This commences proofreading in step-2. Excision of the nucleotide in step-3 resets the system to its initial state.

The schemes in Fig. 1(B,D) show the aa-tRNA selection process by ribosome during translation [29]. Here, E denotes the *E. coli* ribosome with mRNA. Cognate (near-cognate) aa-tRNAs in ternary complex with elongation factor Tu (EFTu) and GTP bind with ribosome in step-1 to form ER (EW). GTP hydrolysis in step-2 results in the complex ER^*^ (EW^*^). The latter can take one of two routes. It can progress to the product P_R_ (P_W_) with the elongation of the peptide chain by one amino acid. Alternatively, it can dissociate in the proofreading step (step-3), rejecting the aa-tRNA.

In both schemes, we take the rate constants of the W cycle to be related to those of the R cycle through *k*_±*i,W*_ = *f*_±*i*_*k*_±*i,R*_*, i* = 1, 2, 3 and similarly, for the catalytic step, *k*_*p,W*_ = *f*_*p*_*k*_*p,R*_. The set of rate constants *k*_1*,R/W*_*, k*_−3*,R/W*_ are effectively first-order containing the substrate concentrations. The factors *f*_*i*_ provide the energetic discrimination between the R and W pathways. Completion of one cycle (returning to the starting state E) effectively amounts to hydrolysis of one dNTP molecule for DNA replication and one GTP molecule for aa-tRNA selection. The chemical potential difference, Δ*µ* (in units of *k*_B_*T*) is equal for both the cycles [19]

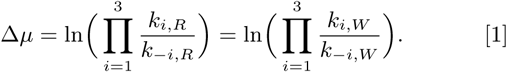

This leads to the condition

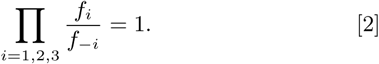

### Accuracy and speed from first-passage description

We determine the error and speed of the substrate selection kinetics from the first-passage probability density [27, 28]. With this method we can analyze an arbitrary catalytic reaction scheme and focus on the transitions starting from the initial state *E* that lead to the final state P_R_. The description allows us to get analytical expression for both speed and accuracy for an arbitrary set of kinetic parameters. Therefore, we allow different rates for the right and wrong substrates for each step of the network as experimentally observed. Furthermore, we do not assume any step to be completely irreversible. Otherwise, the chemical potential difference over the cycle would diverge (see Eq.(1)). This difference is linked to the hydrolysis of some energy-rich molecules supplying large but finite free energy.

Let us denote *F*_R,E_(*t*) as the probability density to reach state P_R_ at time *t* for the first time before reaching state P_W_ if the system is in state E at time *t* = 0. The corresponding probability density *F*_W,E_(*t*) is specified in the same manner. The evolution equations of *F*_R*/*W,E_(*t*) are known as the backward master equation [27, 30]. It is more convenient to solve them in Laplace space (see SI). We define the error, *η* as the ratio of the probabilities to reach the end states P_R/W_ (also called the splitting probability [27]) given by

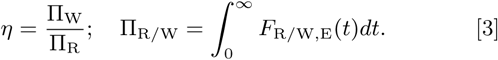

It is important to note that this definition is equivalent to the traditional[9, 21] one defined as the ratio of the wrong product formation rate to the right one (See SI).

The speed of a reaction is naturally quantified by the net rate of the product formation. As any chemical reaction rate, it can be defined as the inverse of the mean first-passage time (MFPT), *i.e.*, the mean time it takes to cross the energy barrier that separates reactants and products for the first time. For example, a well-known application of this approach for single-molecule MM kinetics results in the traditional expression for the rate as the inverse of the MFPT [31]. We note that the speed towards the correct product can nevertheless be affected by the presence of the incorrect substrate. Thus it is important to consider them together in contrast to the prevalent measure of the speed in literature neglecting the presence of the wrong pathway [21]. In our case, the expression of the mean first passage time to reach each product state is given by the first moment of the corresponding probability density [27]

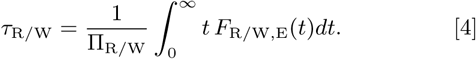

In what follows, we focus on *τ*
_R_ as the measure of speed and denote it simply by *τ*.

## Results

While our formalism can be applied to an arbitrary KPR scheme, we’ve chosen to study two fundamentally important biologically processes — DNA replication and translation. These processes are best characterized in terms of underlying kinetic parameters and we can study the speed-accuracy trade-off in the biologically relevant parameter region. Notably, despite differences in parameters and KPR mechanisms for the two case studies we reach similar conclusions for both.

### Importance of speed over accuracy in DNA replication by T7 DNA polymerase

The T7 DNAP enzyme catalyzes the polymerization of a DNA primer over a template strand [14]. Wrongly incorporated dNTP is removed by the proofreading mechanism that involves the exonuclease site of DNAP [23]. The model parameters of the corresponding reaction network (Fig. 1(A)) are listed in Table S1 (see SI). They are based on the experimental data of Wong *et al.* [23]. We do not consider dissociation of the DNA from the enzyme in our model. This is justified due to the faster polymerization rates in the R path and higher exonucleolytic sliding rate in the W path (see Table S1).

The error, *η* varies among three limits as a function of the polymerization rate constant, *k*_1*,R*_(= *k*_*p,R*_) [23] with fixed *f*_1_*, f*_*p*_. All of them are lower bounds obtained in the limit *k*_1*,R*_ → 0 (*η*_L_), *k*_1*,R*_ → ∞ (*η*_H_) and an intermediate case with *k*_1*,R*_. *k*_−1*,R*_ (*η*_M_). Explicit expressions follow from the general one for *η* (see SI). Here, we give suitable ratios of these limits to understand the error variation pattern

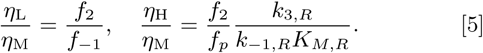

Here, 
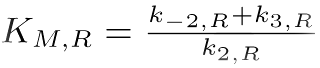
. From the experimental parameter values (see Table S1) and Eq.(5), we expect *η*_M_*< η*_L_*, η*_H_. In other words, the system has a minimum error at some intermediate polymerization rate. This is indeed the case as shown in Fig. 2A. On the other hand, the MFPT, *τ* decreases, and hence the speed increases, monotonously with increase in *k*_1*,R*_ (see Fig. 2B). The range of *τ* also spans several orders of magnitude. The *η*–*τ* curve is shown in Fig. 2C. Negative slope of this curve indicates speed-accuracy trade-off, *i.e.*, higher accuracy (lower *η*) corresponds to lower speed (higher *τ*) and vice versa. It is evident from Fig. 2C that there is a trade-off *only when the polymerization rate constant becomes greater than the value corresponding to the minimum error*. We call this branch with negative slope the trade-off branch. For lower values of *k*_1*,R*_, error and MFPT change in the same direction. This branch with positive slope of the *η*–*τ* curve is denoted as the non-trade-off branch. Intuitively, the lack of trade-off for low polymerization rates arises due to different magnitudes of these rates between the right and wrong pathways; the latter has much smaller rates. When the polymerization rate is sufficiently smaller than the Pol-Exo sliding rate, correct substrate incorporation must undergo lots of unnecessary proofreading cycles. These futile cycles adversely affect the right pathway more, thereby compromising both speed and accuracy. Thus, *the speed-accuracy trade-off might not be always observed* during the proofreading processes, and this is our first important result.

**Fig. 2.**
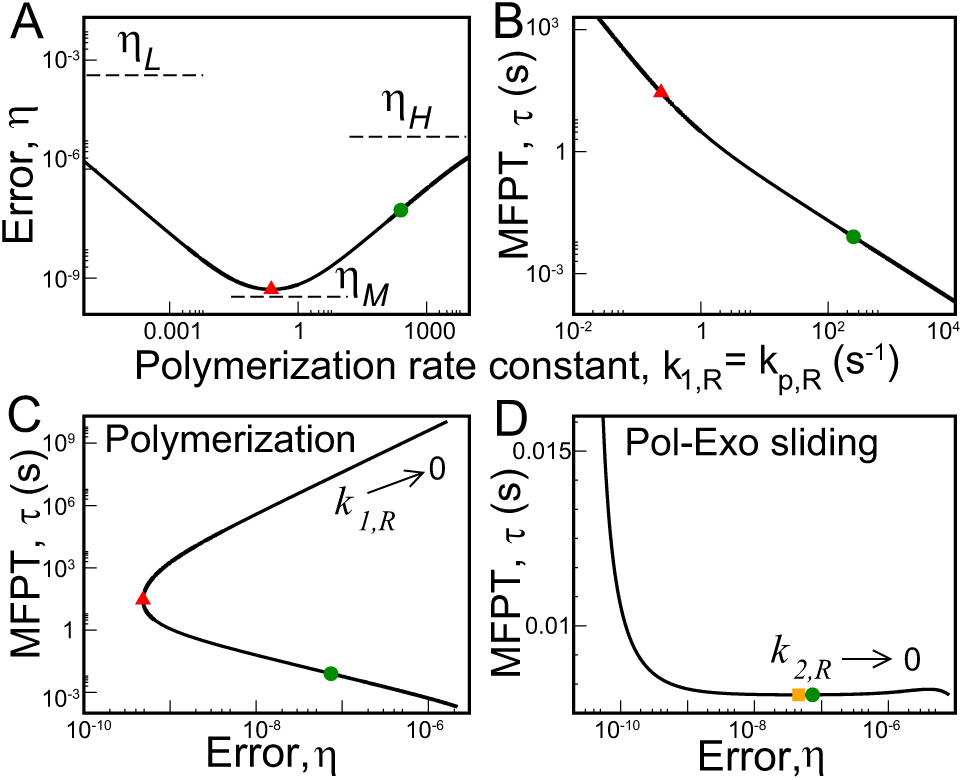
Speed-accuracy trade-off for T7 DNA polymerase. (A) The change in error *_*, as a function of the polymerization rate constant *k*_1*,R*_ (= *k*_*p,R*_). The error is bounded by the predicted limits, Eq.(5). The green circle indicates the position of the actual system that is far away from the minimum error (red triangle). (B) Variation of MFPT, *τ* with *k*_1*,R*_. The red triangle gives the *τ* value corresponding to the minimum in *η*. (C) *η*–*τ* curve for the polymerization step. (D) *η*–*τ* curve for the Pol-Exo sliding step involved in proofreading generated by varying *k*_2*,R*_ keeping *f*_2_ fixed (semi-log plot). There is a local minimum in *τ* (yellow square) near the actual value (green dot).

The actual system (green circle) is situated on the trade-off branch of the *η*–*τ* curve in Fig. 2C. It lies far away from the minimum error point (red triangle). In particular, the minimum error is ~ 150-fold lower than that of the actual system. However, to achieve this minimum error, the system’s speed would drop by ~ 3500-fold. Thus, *the polymerization rate constant is selected to achieve high-enough speed*. Significant amount of accuracy is lost in the process. The system can further lower the MFPT by moving down the slope of the *η*–*τ* curve. But, that means giving up more accuracy. So, of course, there is also a tolerable upper level of *η*.

The Pol-Exo sliding is an important step in error correction. The *η*–*τ* curve for this step is plotted in Fig. 2D. The minimum error value is approached in an asymptotic fashion at very large *k*_2*,R*_. In contrast, the global minimum of MFPT is obtained in the *k*_2*,R*_ → 0 limit. The MFPT also has a local minimum (yellow square) and a local maximum at finite *k*_2*,R*_. Interestingly, *the actual system lies pretty close to this (local)minimum.* In particular, the system’s *τ* value is almost identical to the minimum *τ* (within a less than 0.01%). On the other hand, the corresponding error *η* is ~ 1.6-fold higher than that corresponding to the minimum MFPT. The speed-accuracy trade-off appears after *k*_2*,R*_ crosses the value corresponding to the local minimum in *τ* (and also before the local maximum). So, the system is positioned on the non-trade-off branch of the *η*–*τ* curve. As one moves in either direction from the minimum *τ* point, error can change greatly with slight alteration in *τ* until *η* is too low. Therefore, *speed appears to be more* where *α* = *k*_*p,W*_*/k*_3*,R*_. For the parameter set of the system *important as long as the system remains reasonably accurate*. (see Table S2, *f*_−2_ = 1), one gets *η*_M_*< η*_L_*, η*_H_. Thus, the However, the system can gain lower errors at similar speeds error vs hydrolysis rate curve passes through a minimum. by moving left to the trade-off branch of the *η*–*τ* curve. Then, Interestingly, there is also a minimum in *τ* as shown in Fig. what is the reason for not taking that route? We note, that 3A. The two minima are at different *k*_2*,R*_ values though. The the proofreading pathway resets the system to the starting speed-accuracy trade-off occurs between the minimum *η* (red n condition without progressing to product formation. There triangle, WT) and the minimum *τ* (yellow square, WT) points. fore, speed-up in proofreading rate can increase the associated As is evident from Fig. 3A, all the systems are positioned close extra energy cost [11]. This may restrict the system to go to to the minimum *τ* and far away from minimum *η*. For example, the more advantageous regime that has greater KPR rate and so, somewhat larger cost. We will further elaborate on this point in our next case study.

### tRNA selection by *E. coli* ribosome is optimized for speed rather than accuracy with a cost constraint

During translation, the ribosome decodes the mRNA sequences by selecting aa-tRNAs in ternary complex with elongation factor Tu (EFTu) and GTP [4, 15]. Non-cognate aa-tRNAs are removed by proofreading dissociation of the complex from the ribosome A-site after GTP hydrolysis [24, 29]. The model parameters of the network (Fig. 1B) for WT E. coli ribosome are listed in Table S2 (see SI). They are based on the experimental data of Zaher *et al.* [29]. We chose *k*−2*, R* = *k*−3*, R* = 10−3 *s*−1 to ensure that both step-2 and step-3 are nearly irreversible [29]. There remains one free parameter *f*−2 (as *f*−3 gets fixed from Eq.(2)). We assumed *f*−2 = 1 but our main conclusions are independent of this choice (see SI, Fig. S1).

We show the *η*–*τ* curves for GTP hydrolysis and ternary complex binding steps in Fig. 3A and 3B, respectively, for three varieties of E. coli ribosome. One is the wild-type (WT) and the other two are mutants. One mutant, *rpsL141*, is hyper-accurate (HYP) and the other mutant, *rpsD12*, is more error-prone (ERR) than WT [29]. Variation of the hydrolysis rate constant, *k*2*, R* (keeping *f*2 fixed) results in quite large changes in error and MFPT. The trends are similar for all the three systems (see Tables S3, S4 in SI for parameter sets of mutants). As for the polymerization steps in DNA replication, error varies among three bounds for the GTP hydrolysis step. They are obtained from the general expression of *_* (see SI) for low, intermediate and high values of *k*2*, R*

**Fig. 3.**
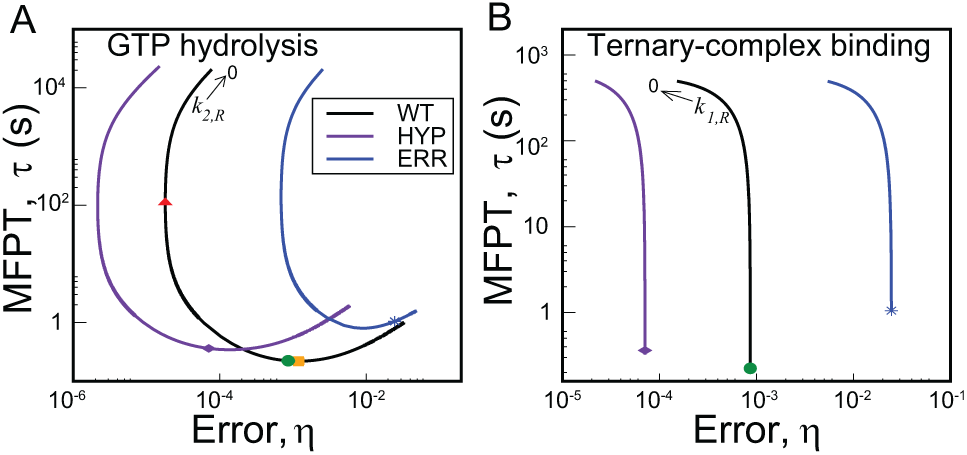
Speed-accuracy trade-off in aa-tRNA selection by three varieties of *E. coli*ribosome. One is the wild-type (WT). The other two are hyper-accurate (HYP) and more error-prone (ERR) mutants. (A) *η*–*τ* curves for the GTP hydrolysis step. The actual system (green circle, WT) is situated close to the minimum *τ* (yellow square) and far away from minimum *η* (red triangle). This is also true for the mutants. (B) Speed-accuracy trade-off for the ternary complex binding step. The minimum error is achieved in the *k*_1*,R*_ → 0 limit. MFPTs for all the systems are close to saturation.

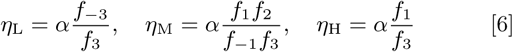

Where *α* = *k^p^, W /k*_3_*, R*. For the parameter set of the system (see Table S2, *f*−2 = 1), one gets *η*M <*η*L*η*H. Thus, the error vs hydrolysis rate curve passes through a minimum. Interestingly, there is also a minimum in *τ* as shown in Fig. 3A. The two minima are at different *k*2*, R* values though. The speed-accuracy trade-off occurs between the minimum *η* (red triangle, WT) and the minimum *τ* (yellow square, WT) points. As is evident from Fig. 3A, all the systems are positioned close to the minimum *τ* and far away from minimum *η*. For example, the WT ribosome would become ~500-fold slower to achieve the minimum error although the latter is ~50-fold lower than the actual value. Hence, *speed is preferred to accuracy*. We tested the generality of this claim against multiple parameter variations (see SI, Fig. S2) and it appears that speed is indeed more important. The robustness of this result is also tested successfully against fluctuations of the rate constants (see Fig. S3). Interestingly, the WT ribosome is faster and hence, better optimized for speed than both the mutants. It is important to note that the more accurate mutant HYP was not chosen by the nature. This point further emphasizes the importance of speed over accuracy in translation.

The change in the ternary complex binding rate constant, *k*_1, *R*_ (with *f*1 fixed) also affects both the error and the MFPT significantly but on a smaller scale than hydrolysis (see Fig. 3B). There is always a trade-off between speed and accuracy unlike the cases studied so far. The minimum in error is obtained in the *k*_1, *R*_ → 0 limit whereas the maximum speed is achieved for very large *k*_1, *R*_. With increase in *k*_1, *R*_, *τ* falls several orders in magnitude. Interestingly, all the systems have *τ* values almost identical to their respective saturation limits. To attain that, they sacrifice an order of magnitude in terms of accuracy. Therefore, regarding speed-accuracy trade-off, the system is inclined to be faster with higher but tolerable error.

Next, we explore the effects of variation of the proofreading step rate constant, *k*_3, *R*_ (keeping *f*_3_ fixed) on system performance. The resulting *η*–*τ* diagram is plotted in Fig. 4A for the WT ribosome. The global minimum of *τ* is obtained in the *k*_3, *R*_ → 0 limit whereas, the minimum of *η* lies in the large *k*_3, *R*_ limit. There is a local minimum (yellow square) of *τ* at some intermediate *k*_3_, *R* along with a local maximum. Mutant ribosomes have similar trends (not shown in figure). The nature of the *η*–*τ* curve is qualitatively similar to that obtained for the proofreading Pol-Exo sliding step in DNA replication (see Fig. 2D) with two speed-accuracy trade-off branches. The actual system (green circle) is located on the non-trade-off branch that links the two trade-off branches of the *η*–*τ* curve. It has a MFPT close to (~1.1-fold higher) the (local) minimum *τ* value. However, the minimum *τ* point also has a ~5-fold lower *η*. More important is the fact that, the system can attain much lower errors with similar, even slightly higher, speeds if it moves left up to a certain level (the dashed line in Fig. 4A). So, what prevents the system to gain both in speed and accuracy?

**Fig. 4.**
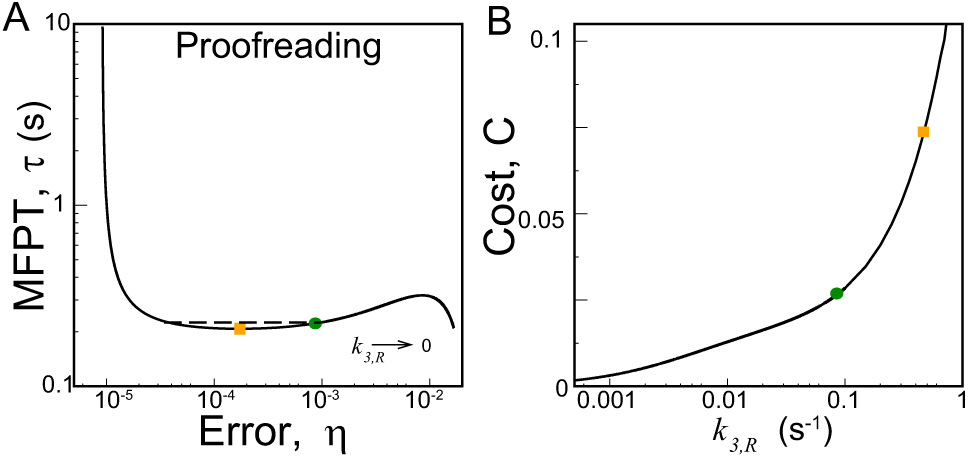
(A)*η*–*τ* diagram for the proofreading step. The actual system (green dot,WT) has a *τ* value similar to the local minimum in *τ* (yellow square). The dashed line shows the range up to which the system can lower the error with no loss in speed. (B) The proofreading cost, C as a function of *k*_3*,R*_. It’s value at the local minimum in *τ* (yellow square) is ~ 3-fold higher than that of the actual system (green dot).

Since correction by proofreading resets the system without a product formation it has a cost associated with futile cycles where the correct substrate was inserted and then removed. The cost of proofreading, *C* is defined as the ratio of the resetting flux to the product formation flux including both R and W pathways [11] (see SI). This gives a measure of the amount of extra energy-rich molecules consumed due to the presence of the proofreading step. Specifically, the cost C can quantify the moles of dNTP(or GTP) used for proofreading per mole of product [11]. This quantity can be easily computed from our formalism and investigated as a function of kinetic parameters. In particular, we quantify how the cost of proofreading changes with the increase in *k*_3*,R*_ near local minimum of the *η*–*τ* curve in Fig. 4A. The results, shown in Fig. 4B, demonstrate that the cost associated with the (local) minimum *τ* point is ~ 3-fold higher than that of the actual system. This cost-disadvantage (3-fold higher GTP consumption per amino acid!) may restrict the system from gaining the available advantage in both speed and accuracy. Similar consideration may also be responsible for the nature of trade-off exhibited in the DNA replication case (Fig. 2D).

## Discussion

Evolution has optimized the kinetic parameters of biological enzymes to achieve the desired levels of accuracy and speed at various stages of biological information flow. In this study, by examining how the balance between speed and accuracy changes with variation of the underlying kinetic parameters, we gain new insights of the important priorities for this optimization. To this end, we focus on two fundamental examples of biological proofreading networks: DNA replication and protein translation. In both the cases, the systems tend to achieve maximum speed by losing significant accuracy. However, the speed-accuracy trade-off only occurs in the limited region of the parameter space, e.g., after the polymerization rate in replication passes the minimum error point. In case of translation, the trade-off appears between the minima in error and the MFPT for the GTP hydrolysis step. Similar conclusion about the importance of speed over accuracy is reached by varying the rates of the proofreading steps in both systems. Although higher proofreading rates can further improve the accuracy without losing much in speed, the associated energy cost of proofreading may restrict further improvements on an already acceptable speed and accuracy.

An important new insight from the above analyses is that, speed-accuracy trade-off is not universally present and its occurrence depends on the specific values of kinetic rates. Biologically that implies that mutations or application of drugs that reduce the enzyme’s accuracy do not necessarily increase its speed and *vice versa*. The widespread view of a compromise between accuracy and speed is mainly based on their dependence on the effective catalytic rate of the process [9, 21]. Indeed, the larger catalytic rate – the higher speed and the lower accuracy. However, the role of other steps, like hydrolysis and proofreading, are not as straightforward. Our study reveals that, for these steps, trade-offs are present only over a certain range of rates. The partitioning of the error-time curves into trade-off and non-trade-off branches clarifies the distinct roles of various transitions and the molecular mechanisms of the speed-accuracy optimization. Our conclusions are also supported by a more advanced analysis of the maximum speed vs accuracy curves using Pareto fronts, as explained in detail in the SI.

The analysis of speed-accuracy trade-off for different mutant varieties of *E. coli* ribosome further confirms the importance of speed over accuracy. The WT and two mutants (HYP and ERR) lie close to the minimum MFPT point on the error-time curves (Fig.3A). However, the WT and HYP ribosomes are on the trade-off branch whereas, the ERR mutant is on the non-trade-off branch. Thus, movement down the slope towards the trade-off branch would raise both accuracy and speed for the ERR ribosome. That is how the WT ribosome may have evolved from the more erroneous ERR type. However, any further movement upwards along the trade-off branch means slowdown with a lower error. This leads to the more accurate (HYP) mutant. Rejection of the latter as the natural choice implies that optimization of speed is critical. We note that comparison of *E. coli* growth rates with WT and mutant ribosomes already indicates such an optimization [21, 32]. However, according to the prevailing notion on the ever-present compromise between error and speed, the more erroneous (ERR) ribosome should be faster. Hence, the hindered growth for ERR mutant was ascribed presumably to less-active proteins [33]. Our results indicate that, not only the accuracy but also the speed of peptide-chain elongation can be smaller for the ERR mutant.

Despite different schemes and parameter values of the replication and translation networks, there appears to be a general mechanism of error correction. This becomes apparent from the trade-off diagrams for the proofreading step. Rate constant of the proofreading step in both the cases is selected such that speed of the system is close to the maximum-possible one. The actual systems reside on the non-trade-off branch of their respective error-time curves. Biologically that implies that mutation that slightly speeds-up the proofreading step would lead to increase in both speed and accuracy of the enzymes. However, we show that such mutation would also increase energetic costs of proofreading. This extra cost does not allow the systems to further reduce the error and MFPT. Furthermore, the most interesting feature for both the systems is the proximity of the MFPT value to the local minimum which is similar in magnitude to the global minimum. Hence, for both case studies the KPR rate is fine-tuned so that the loss in speed is insignificant compared to the improvement in accuracy.

Our results on the accuracy-speed trade-off in two important biological networks reveal similar strategies to optimize these two vital quantities. Rates of the steps like substrate binding, hydrolysis (of intermediates) and catalysis seem to be chosen to enhance speed at the cost of accuracy. On the other hand, proofreading or error-correction steps seem to be selected to have such rates that the error is reduced sufficiently with almost no loss in speed. Therefore, between the maximization of accuracy and speed, biological systems appear to give precedence to the latter. Tolerable levels of error and cost of error-correction act as constraints to tailor the speed. It is interesting to note here that experimentally observed distribution of discriminatory steps is not optimal from the point of view of minimizing error[34]. For example, for ribosome the rates of the catalytic step are significantly different between the incorporation of the right and wrong amino-acid in the polypeptide chain. While this may be suboptimal in terms of error minimization [34], it allows for the proofreading rate to be much faster than catalytic rate for a wrong substrate and much slower than the catalytic rate for the right substrate. As a result, ribosome avoids futile cycles (correcting the errors it did not make) improving speed and energy cost. This observation gives additional support to our arguments that biological systems distribute discrimination to better optimize speed and not accuracy (see SI). Our study, thus, presents a coherent quantitative picture of how the ultimate balance between accuracy and speed is achieved by adjusting various rates in distinct ways. It will be important to test our predictions in other systems and organisms. We believe, this will further help to elucidate the fundamental mechanisms of proofreading processes in biological systems.

## ACKNOWLEDGMENTS

This work is supported by Center for Theoretical Biological Physics NSF Grant PHY-1427654. ABK also acknowledges the support from the Welch Foundation (Grant C-1559) and from the NSF (Grant CHE-1360979).

## Supporting Information

Banerjee et al. 10.1073/pnas. XXXXXXXXXX

### SI Text

#### First-passage probability density: Evolution equations

We study the kinetics of the proofreading networks in terms of the first-passage process. The key quantity to characterize the dynamic properties of the system is the first-passage probability density. Let us denote *F*_R,i_(*t*) as the first-passage probability density to reach the right (R) end-state at time *t* forming product P_R_ for the first time before reaching the wrong (W) end-state (product P_W_), starting from state *i* at time *t* = 0 (see Fig. 1). The corresponding probability density *F*_W,i_(*t*) is defined in the same manner. The equations describing the time-evolution of the first-passage probability densities are generally known as the backward master equations. To solve them, we introduce **F**_**R**_ = (*F*_R,E_, *F*_R,ER_, *F*_R,ER*_,*F*_R,EW_, *F*_R,EW*_)^T^. **F**_**W**_ is constructed in similar fashion.

For the DNA replication network (Fig. 1C), we can write

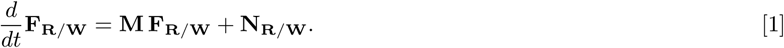

Here, **N**_**R**_ = (0, *k*_*p,R*_*δ*(*t*), 0, 0, 0)^T^ and **N**_**W**_ = (0, 0, 0, *k*_*p,W*_*δ*(*t*), 0)^T^, *δ*(*t*) being the Dirac-delta function. The transition matrix

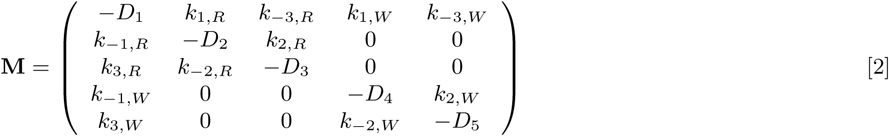

where *D*_1_ = (*k*_1,*R*_ + *k*_1,*W*_ + *k*_−3,*R*_ + *k*_−3,*W*_), *D*_2_ = (*k*_−1,*R*_ + *k*_2,*R*_ + *k*_*p*,*R*_), *D*_3_ = (*k*_−2,*R*_ + *k*_3,*R*_), *D*_4_ = (*k*_−1,*W*_ + *k*_2,*W*_ + *k*_*p*,*W*_), *D*_5_ = (*k*_−2,*W*_ + *k*_3,*W*_).

It is more convenient to solve Eq.(1) in Laplace space. The Laplace transform of *F*_R*/*W,i_(*t*) is defined as

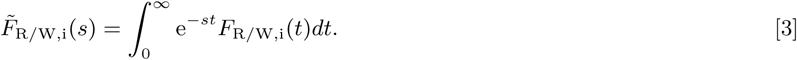

Under this transformation, with the initial condition *F*_R*/*W,i_(*t* = 0) = 0, (*i* ≠ *R/W*), Eq.(1) becomes

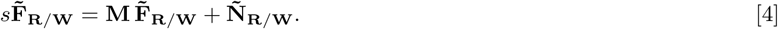

Here, 
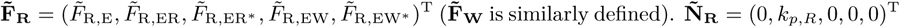
 and 
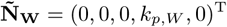
. From the algebraic set of equations, Eq.(4), one gets the desired quantities 
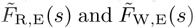
, *i.e.*, the first-passage probability density to reach one of the two end-states for the first time before reaching the other, starting from state-E. The corresponding splitting probabilities to reach either of the end-states are expressed as

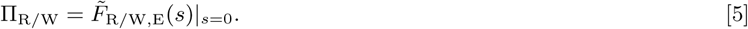

We employ the same methodology for the translation network in Fig. 1D. Note that now the end-states are linked with the states ES^*^ instead of ES as in replication. The transition matrix **M** has similar structure but now with *D*_2_ = (*k*_−1*,R*_ + *k*_2*,R*_), *D*_3_ = (*k*_−2*,R*_ + *k*_3*,R*_ + *k*_*p,R*_), *D*_4_ = (*k*_−1*,W*_ + *k*_2*,W*_), *D*_5_ = (*k*_−2*,W*_ + *k*_3*,W*_ + *k*_*p,W*_). We also get **N**_**R**_ = (0, 0, *k*_*p,R*_ *δ*(*t*), 0, 0)^T^ and **N**_**W**_ = (0, 0, 0, 0, *k*_*p,W*_*δ*(*t*)). The corresponding Laplace transforms are 
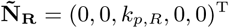
 and 
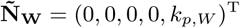
.

#### Error in replication and translation

The error is defined as the ratio of the splitting probabilities

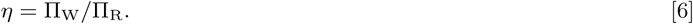

The exact expression of error for the DNA replication network (Fig. 1C in main text) is given by

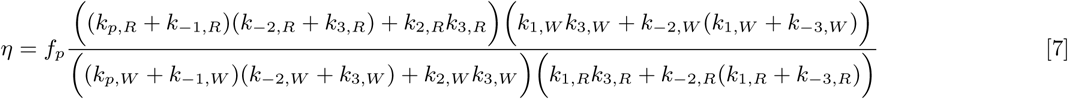

The experimental data suggest the following limits: *k*_−1*,S*_*<< k*_*p,S*_, *k*_−3*,S*_*<< k*_1*,S*_ (S = R or W). Under these limits, one gets

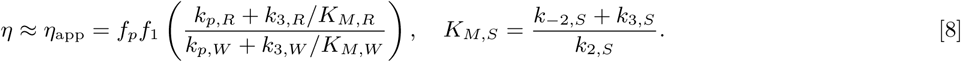

It is easy to see that the translation network (Fig. 1D in main text) is related to the replication network by the following transformation of rate constant indices: 
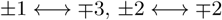
. Then, from Eq.(7), the general expression of error for the translation network comes out as

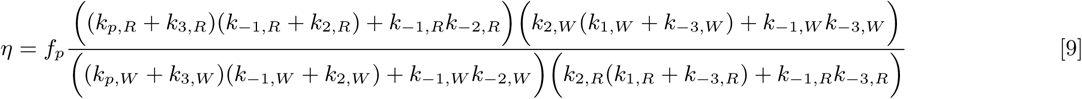

If one considers no discrimination in step 2 and the catalysis step, *i.e.*, sets *f*_2_ = 1 = *f*_−2_ = *f*_*p*_, then Eq.(9) reduces to the form as originally derived by Hopfield [Hopfield JJ (1974) Proc Natl Acad Sci USA 71:4135–4139, Eq.(5)]. This clearly shows the validity of our approach. According to experimental data, GTP hydrolysis (step-2) and proofreading (step-3) steps are mainly unidirectional. Then, taking the limits *k*_−2*,S*_, *k*_−3*,S*_ → 0, we get the approximate form of error as

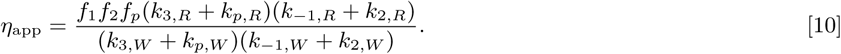

#### Dependence of minimum error on the ratio of proofreading rate to catalysis rate

The minimum in error is obtained when the system is away from equilibrium but the initial enzyme-substrate binding equilibrium is minimally disturbed. This means *k*_2*,S*_*<< k*_−1*,S*_ (S = R or W) with the magnitude of *k*_2*,S*_ being high-enough to provide sufficient driving chemical potential difference over the cycles. From Eq.(10), we get the minimum error as

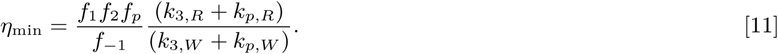

We now consider three different scenarios of *k*_3*,S*_*/k*_*p,S*_.

Case I. 
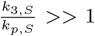
, *i.e.*, the catalysis rates for both right and wrong pathways are small compared to proofreading. From Eq.(11), one gets

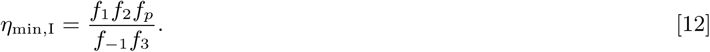

It is satisfactory to see that, if one sets *f*_1_ = 1 = *f*_2_ = *f*_*p*_ and *f*_−1_ = *f*_0_ = *f*_3_, then one achieves the Hopfield limit of minimum error 
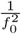
.

Case II. *k*_3*,R*_*<< k*_*p,R*_, *k*_3*,W*_*>> k*_*p,W*_, *i.e.*, the catalysis rate of right pathway dominates proofreading but the opposite is true for the wrong pathway. The WT ribosome as well as the mutants are examples of such cases (see Tables S3-S5). In this case, we have

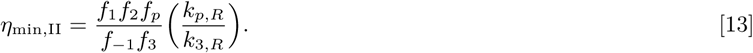

Case III. 
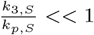
 1, *i.e.*, the catalysis rates for both right and wrong pathways are large compared to proofreading. Then, from Eq.(11) it follows

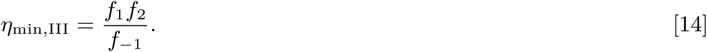

Comparing Eqs.(12)-(14), we have

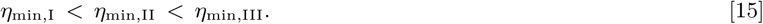

For the sake of completeness, we also determine the minimum error in the DNA replication case. From Eq.(8), large *k*_2*,S*_ gives this limit as

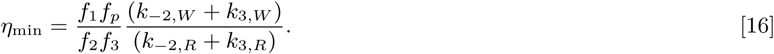

We take *k*_−2*,R*_ = *k*_−2*,W*_, *k*_3*,R*_ = *k*_3*,W*_ (see Table S2) and then 
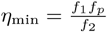
. A high excision (by hydrolysis) rate compared to the Exo-Pol back sliding rate, *i.e.*, *k*_3*,S*_*>> k*_−2*,S*_ gives the same minimum error. On the other hand, *k*_3*,S*_*<< k*_−2*,S*_ gives 
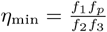
.

#### The conditional mean-first-passage time

The conditional mean first-passage times (MFPTs) to reach the respective end-states (in presence of the other) are defined in terms of the first-passage probability densities in Laplace space as

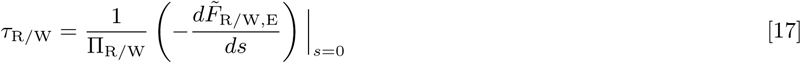

For our purpose, the quantity of interest is *τ*
_R_, the MFPT (’conditional’ term implied) to reach the R end. We denote it simply by *τ*. One can obtain analytical expressions for the MFPT for both the networks using Eq.(17). However, the exact forms come out to be quite unwieldy. To get some theoretical insights, we provide here approximate expressions. These are valid over the relevant parameter ranges, tested for both the networks. We emphasize that all the plots of the MFPTs are generated using the exact expressions.

In case of DNA replication, experimental data indicate *k*_2*,R*_*<< k*_*p,R*_, (1 + *η*) ≈ 1. Then, with the limits *k*_−1*,S*_, *k*_−3*,S*_ → 0, we get

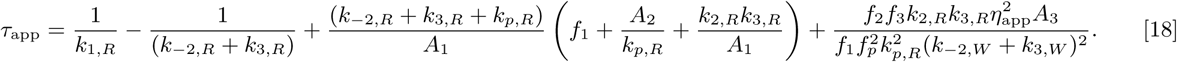

Here, *A*_1_ = *k*_*p*,*R*_(*k*_−2,*R*_ + *k*_3,*R*_), 
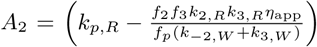
, *A*_3_ = (*k*_*p*,*W*_ + *k*_−2,*W*_ + *k*_2,*W*_ + *k*_3,*W*_) and *η*_app_ is given by Eq.(8).

Kinetic data for the translation network show that *k*_*p*,*R*_ ≫ *k*_3,*R*_, *k*_*p*,*W*_ ≪ *k*_3,*W*_, (1 + *η*) ≈ 1. Then, taking the *k*_−2*,S*_, *k*_−3*,S*_ → 0 limit, one has

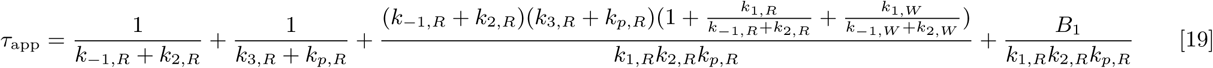

Here, 
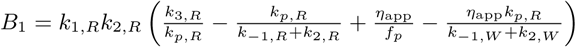
 and *η*_app_ is given by Eq.(10).

#### Cost of proofreading

The enhanced accuracy due to proofreading comes at a price. The KPR step resets the system and prevents it to take the catalytic path. Thus, hydrolysis energy of triphosphate molecules is consumed without product formation. This extra cost or cost of proofreading, C is defined as the ratio of the resetting or proofreading flux to the product-formation flux. Thus,

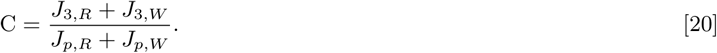

Here, *J*_3*,S*_ = *k*_3*,S*_*P*_ES*_ − *k*_−3*,S*_*P*_E_ (S = R or W) is the flux associated with the proofreading step (step-3) and *J*_*p,S*_ = *k*_*p,S*_*P*_ES*_ is the flux of the catalytic step. *P*_j_ denotes the steady-state probability to find the system in state j. For the DNA replication network, *J*_3*,S*_ = *J*_2*,S*_.

For the aa-tRNA selection network, taking the *k*_−3*,S*_ → 0 limit yields

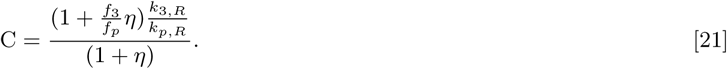

Here, *η* is defined as the ratio of the catalytic fluxes of wrong and right product. A similar simplification, however, does not produce such a compact form for the DNA replication network. We use Eq.(20) to generate the plot in Fig. 4(b) of the main text.

**Fig. S1.**
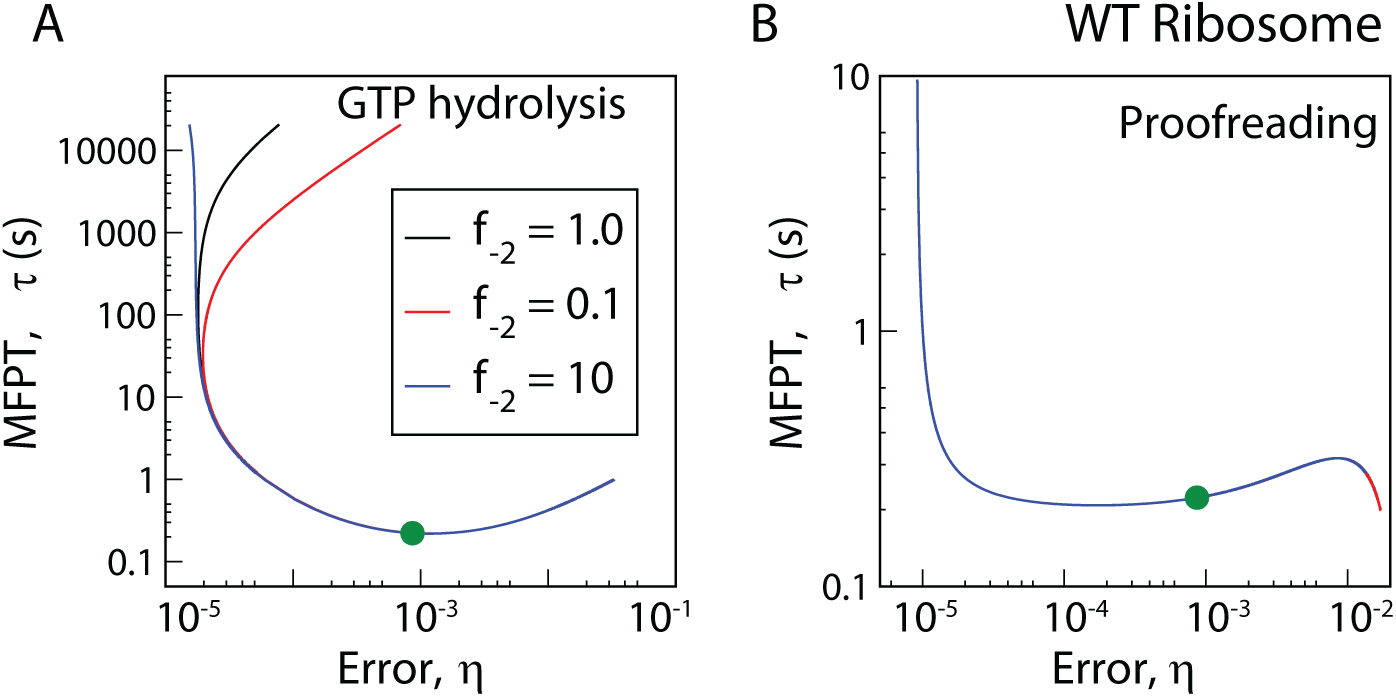
Robustness of error-MFPT trade-off for aa-tRNA selection against change in the parameter*f*_−2_. Trade-off diagrams are generated for the WT ribosome with three different values of the parameter *f*
_−2_ for (A) the GTP hydrolysis step and (B) the proofreading step. The trade-offs remain unchanged near the relevant parameter range for the actual system (green circle, *f*_−2_ = 1.0). For the GTP hydrolysis step with *f*_−2_ = 10, the minimum in error is attained in the *k*_2*,R*_ → 0 limit. However, this qualitative change of trend occurs far away from the region of interest. Therefore, our main conclusion about the importance of speed optimization over accuracy remains valid over relevant parameter ranges.

#### Maximum speed vs accuracy curves and Pareto front

In this section, we employ multiple parameter variation to investigate the question: What is the maximum speed for a given accuracy value for some choice of parameters? To start with, one must note that the global maximum speed available to the system is always the catalytic rate, *k*_*p,R*_, irrespective of the value of other parameters. In other words, if one chooses to vary *k*_*p,R*_ then one can always get a higher speed at higher *k*_*p,R*_. Hence, it is reasonable to first fix the catalytic rate constant. As all the trade-off curves in the main text are determined for fixed energetic discrimination ratios *f*_*i*_, *f*_*p*_, here we also keep them fixed. The reverse rate constants for hydrolysis and proofreading are fixed to low values to ensure the nearly unidirectional nature of these steps.

**Fig. S2.**
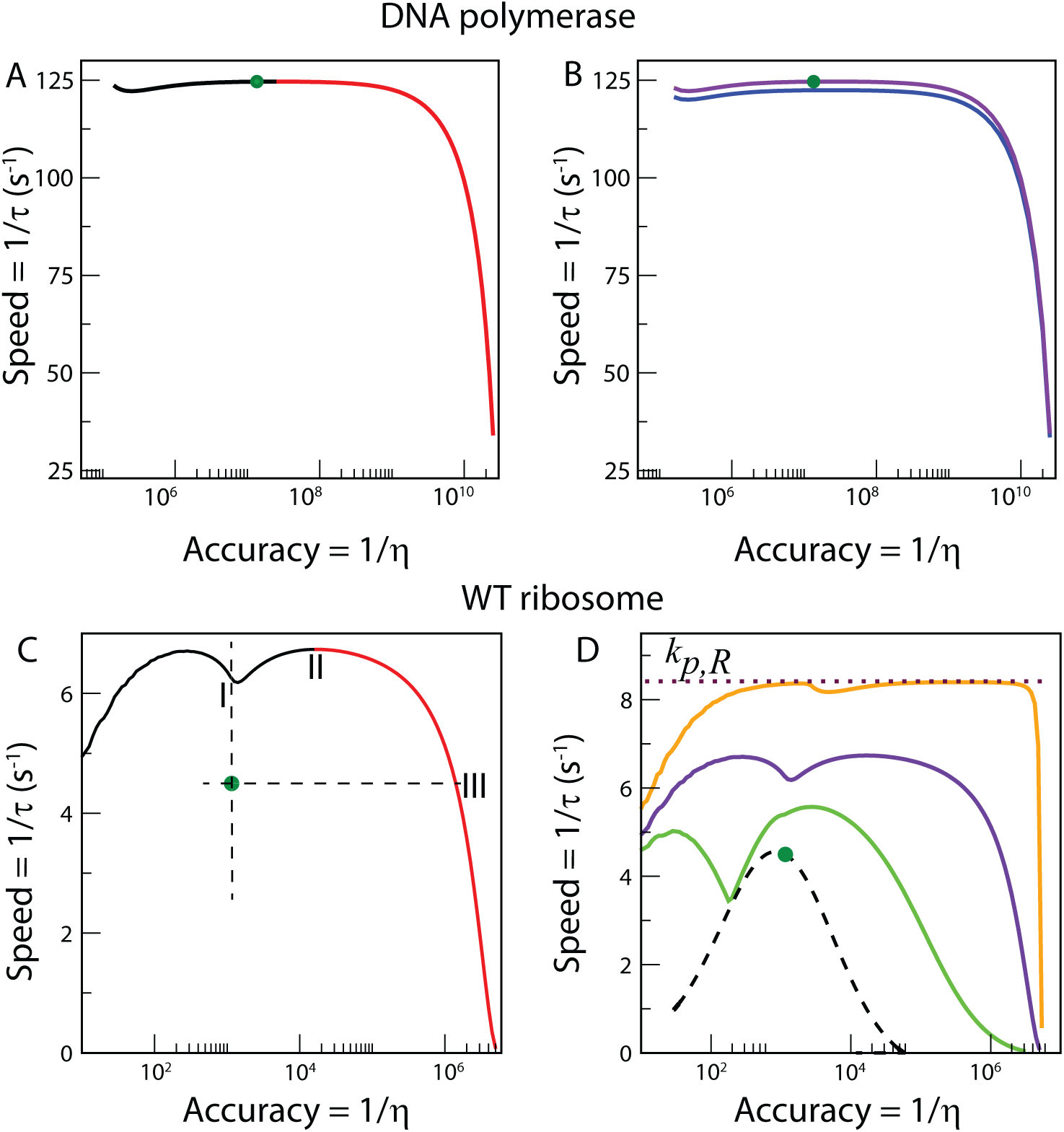
(A) The curve of maximum speed(= 1*/τ*) against accuracy(= 1*/η*) for DNA replication by T7 DNA polymerase. It is generated by varying the rate constants*k*_−2*,R*_, *k*_3*,R*_over a range of10^−6^−10^6^*s*^−1^while *k*_2*,R*_is evaluated at each accuracy value as a function of *k*_−2*,R*_, *k*_3*,R*_, *η* and other fixed parameters. The red portion of the curve shows a Pareto front. The position of the system is indicated with a green dot. (B) Maximum speed vs accuracy curves generated in the same manner described in panel (A). The blue curve is obtained by increasing *k*_−1*,R*_ from 1 *s*^−1^ to 10 *s*^−1^. The violet curve is the same one as in panel (A) and redrawn for the sake of comparison. (C) The curve of maximum speed(= 1*/τ*) against accuracy(= 1*/η*) for aa-tRNA selection WT *E. coli* ribosome. It is obtained by varying the rate constants *k*_2*,R*_, *k*_−1*,R*_ over a range of 10^−6^ − 10^6^*s*^−1^ while *k*_3*,R*_ is evaluated at each accuracy value as a function of *k*_2*,R*_, *k*_−1*,R*_, *η* and other fixed parameters. The red portion of the curve shows a Pareto front. The position of system is indicated with a green dot. (D) Variation of speed with accuracy for different cases of parameter variation for WT ribosome. The curve with dashed black line corresponds to the *η*-*τ* curve in Fig. 3A of main text and the dot on the curve shows the position of the actual system. The other three curves show the variation of the maximum speed with accuracy. The green curve is generated by determining the maximum speed for a given accuracy value by varying the rate constant *k*_2*,R*_ with *k*_3*,R*_ being evaluated as a function of *k*_2*,R*_, *η* and other fixed parameters. The violet curve (the same curve redrawn from panel A) and the orange curve are generated in the same manner described in panel (A). For the orange curve, we set *k*_1*,R*_ = 10^4^ s. The dotted line represents the value of the catalytic rate constant *k*_*p,R*_, the upper boundary of speed.

We determine maximum speed for a given accuracy value by varying different sets of parameter. Some representative curves are shown in Fig. S2 for DNA replication and aa-tRNA selection. In Fig. S2A, the maximum speed vs accuracy curve is generated in the following manner: rate constants *k*_−2*,R*_, *k*_3*,R*_ are varied freely over a range of 10^−6^ − 10^6^*s*^−1^ while *k*_2*,R*
_ is evaluated at each accuracy value as a function of *k*_−2*,R*_, *k*_3*,R*_, *η* and other fixed parameters. The red part of the curve depicts what is known as a Pareto front. By definition, over the Pareto front there is always trade-off between maximum speed and accuracy. All the points over the rest of the curve (and inside) have either lower accuracy or lower speed or both compared to those comprising the Pareto front. It is evident from Fig. S2A that, the actual system lies very close to the maximum speed boundary as well as the Pareto front. The system can increase its accuracy without compromising much in speed by moving onto the Pareto front. However, the cost factor restricts such a strategy similar to the case shown in Fig. 4 of the main text. In Fig. S2B, the blue curve is obtained by increasing *k*_−1*,R*_ from 1 *s*^−1^ to 10 *s*^−1^; the violet curve is the same as in Fig. S2A and redrawn for comparison. As expected, the maximum speed for the blue curve is lower but it is still fairly close to the violet one. On the other hand, decreasing *k*_−1*,R*_ has very little effect on the maximal speed vs accuracy curve and the Pareto front. Hence, the proximity of the system to the maximum speed and the Pareto front is a robust feature.

The maximum speed vs accuracy curves for WT ribosome are shown in Fig. S2C, D. They are generated in the following manner: rate constants *k*_2*,R*_, *k*_−1*,R*_ are varied freely over a range of 10^−6^ − 10^6^*s*^−1^ while *k*_3*,R*_ is evaluated at each accuracy value as a function of *k*_2*,R*_, *k*_−1*,R*_, *η* and other fixed parameters. The actual system lies inside the Pareto front and is relatively close to the maximum speed but not to the maximum accuracy. In Fig. S2D, similar cases of parameter variation are depicted (for details, see caption). The data in Fig. 3A of the main text is also included for comparison. The maximum speed curve goes very close to the upper boundary when we set the rate constant *k*_1*,R*_ to a large value(= 10^4^*s*^−1^, orange curve in Fig. S2D). It is expected as increase in *k*
_1*,R*_ helps to increase the speed as already shown in Fig. 3B of the main text. Thus, even if we were to include *k*_1*,R*_ in our multi-parameter variation, the maximum speed is expected to arise at the upper boundary of the range of *k*_1*,R*_. Now, it is crucial to figure out what are the actual parameter values for some suitably-chosen points on the maximum speed vs accuracy curves. For this purpose, we consider the curve in Fig. S2C and pick three points on the curve as follows: I. with accuracy equal to that of the system; II. the maximum of the maximum speed vs accuracy curve and III. with maximum speed equal to that of the system. The corresponding parameter values are listed in Table S5. It is evident from the data that all three positions are unrealistic for the system to reside on, particularly in the light of experimentally measured kinetic constants (see Table S2). It is important to note that, as the system lies inside the Pareto front, one can not rule out the possibility of improving both speed and accuracy with affordable cost. However, it is not quite clear what additional physical and/or biochemical constraints could keep the system away from the Pareto front.

---

**Fig. S3.**
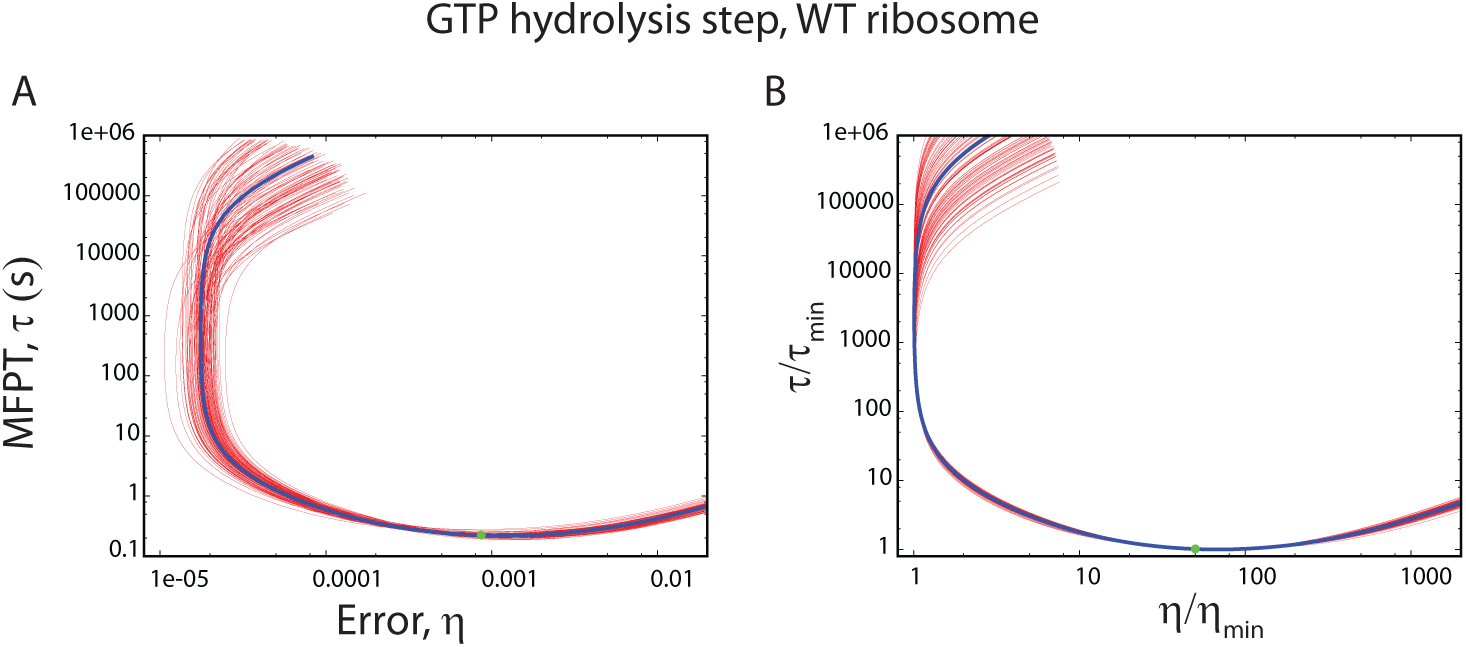
Robustness of the error-MFPT trade-off for the GTP hydrolysis step with respect to random variation of various rate constants. The factors*f*_*i*_, *f*_*p*_are kept fixed. The rate constants *k*_−2*,R*_, *k*_−3*,R*_ are varied in the range 10^−3^ − 10^−5^ by sampling from a log-uniform distribution. Other rate constants are sampled from a Gaussian distribution with the standard deviation equal to the experimental error bars in the kinetic data of Zaher *et al* [Zaher HS, Green R (2010) Molecular Cell 39(1):110–120]. (A) Bunch of trade-off curves generated from random sampling. The blue curve shows the trade-off with no errors and the green circle shows the system’s position. (B) Trade-off curves generated by scaling error and MFPT values of each curve with their respective minimum. It is evident from the plots that the preference of the system to achieve maximum speed rather than maximum accuracy is quite robust against parameter fluctuations.

**Table S1.** Model parameters for DNA replication by T7 DNAP.

| Parameter | Value ($s^{-1}$) | Parameter | Value |
| --- | --- | --- | --- |
| $k_{1,R}$ | 250 | $f_1$ | $8 \times 10^{-6}$ |
| $k_{-1,R}$ | 1 | $f_{-1}$ | $1 \times 10^{-5}$ |
| $k_{2,R}$ | 0.2 | $f_2$ | 11.5 |
| $k_{-2,R}$ | 700 | $f_{-2}$ | 1 |
| $k_{3,R}$ | 900 | $f_3$ | 1 |
| $k_{p,R}$ | 250 | $f_p$ | $4.8 \times 10^{-5}$ |

**Table S2.** Model parameters for aa-tRNA selection by WT *E. coli* ribosome.

| Parameter | Value ($s^{-1}$) | Parameter | Value |
| --- | --- | --- | --- |
| $k_{1,R}$ | 40 | $f_1$ | 0.675 |
| $k_{-1,R}$ | 0.5 | $f_{-1}$ | 94 |
| $k_{2,R}$ | 25 | $f_2$ | $4.8 \times 10^{-2}$ |
| $k_{3,R}$ | $8.5 \times 10^{-2}$ | $f_3$ | 7.9 |
| $k_{p,R}$ | 8.415 | $f_p$ | $4.2 \times 10^{-3}$ |

**Table S3.**
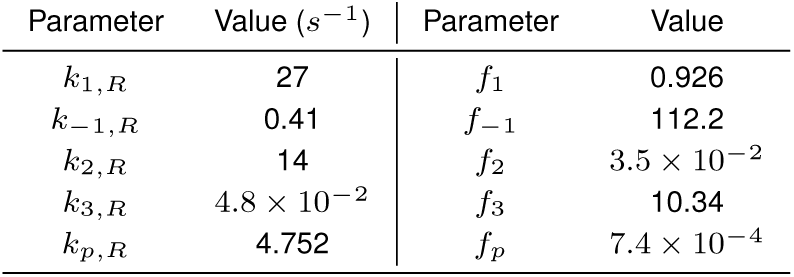
Model parameters for aa-tRNA selection by hyper-accurate (HYP, *rpsL141*) mutant *E. coli* ribosome.

| Parameter | Value ($s^{-1}$) | Parameter | Value |
| --- | --- | --- | --- |
| $k_{1,R}$ | 27 | $f_1$ | 0.926 |
| $k_{-1,R}$ | 0.41 | $f_{-1}$ | 112.2 |
| $k_{2,R}$ | 14 | $f_2$ | $3.5 \times 10^{-2}$ |
| $k_{3,R}$ | $4.8 \times 10^{-2}$ | $f_3$ | 10.34 |
| $k_{p,R}$ | 4.752 | $f_p$ | $7.4 \times 10^{-4}$ |

**Table S4.** Model parameters for aa-tRNA selection by more-erroneous (ERR, *rpsD12*) mutant *E. coli* ribosome.

| Parameter | Value ($s^{-1}$) | Parameter | Value |
| --- | --- | --- | --- |
| $k_{1,R}$ | 37 | $f_1$ | 0.973 |
| $k_{-1,R}$ | 0.43 | $f_{-1}$ | 9.3 |
| $k_{2,R}$ | 31 | $f_2$ | 0.126 |
| $k_{3,R}$ | $7.7 \times 10^{-2}$ | $f_3$ | 7.65 |
| $k_{p,R}$ | 7.623 | $f_p$ | $4.1 \times 10^{-3}$ |

**Table S5.** Parameter values corresponding to the three points on the maximum speed vs accuracy curve in Fig. S2A for WT *E. coli* ribosome. All the rate constants are in *s*^−1^.

| Point | $k_{-1,R}$ | $k_{2,R}$ | $k_{3,R}$ |
| --- | --- | --- | --- |
| I | $5.6 \times 10^5$ | $10^6$ | $3.3 \times 10^{-4}$ |
| II | $6.1 \times 10^5$ | $10^6$ | 0.51 |
| III | $10^6$ | $9.6 \times 10^5$ | 8.7 |

